# Programmable and Dynamic DNA Localisation at Synthetic Cell Membranes

**DOI:** 10.64898/2026.07.13.738173

**Authors:** Chelsea Dack, Bingkun Li, Charlie Newell, Michael J. Booth

## Abstract

Spatial and temporal organisation of membrane-associated components is fundamental to cellular signalling, yet remains difficult to engineer in minimal synthetic systems. In synthetic cells, DNA and RNA nanotechnology offer programmable molecular organisation at membranes, while *in vitro* transcription (IVT) enables gene expression-driven regulation. However, integrating these systems within cell-like compartments, such as giant unilamellar vesicles (GUVs), remains challenging due to undesirable interactions between transcription machinery and nucleic acid assemblies. Here, we present a modular strategy that couples *in situ* RNA production to dynamic DNA localisation at GUV synthetic cell membranes. RNA strands, transcribed within GUVs, function as linkers that recruit DNA-conjugated cargo to lipid membranes, enabling programmable spatial organisation. Using this framework, we achieved reversible membrane localisation through toehold-mediated strand displacement and RNase H-mediated degradation. This work establishes a gene expression-driven platform for programmable and dynamic control of membrane-associated components in synthetic cells, providing a foundation for constructing dynamic signalling assemblies and higher-order cellular behaviours.

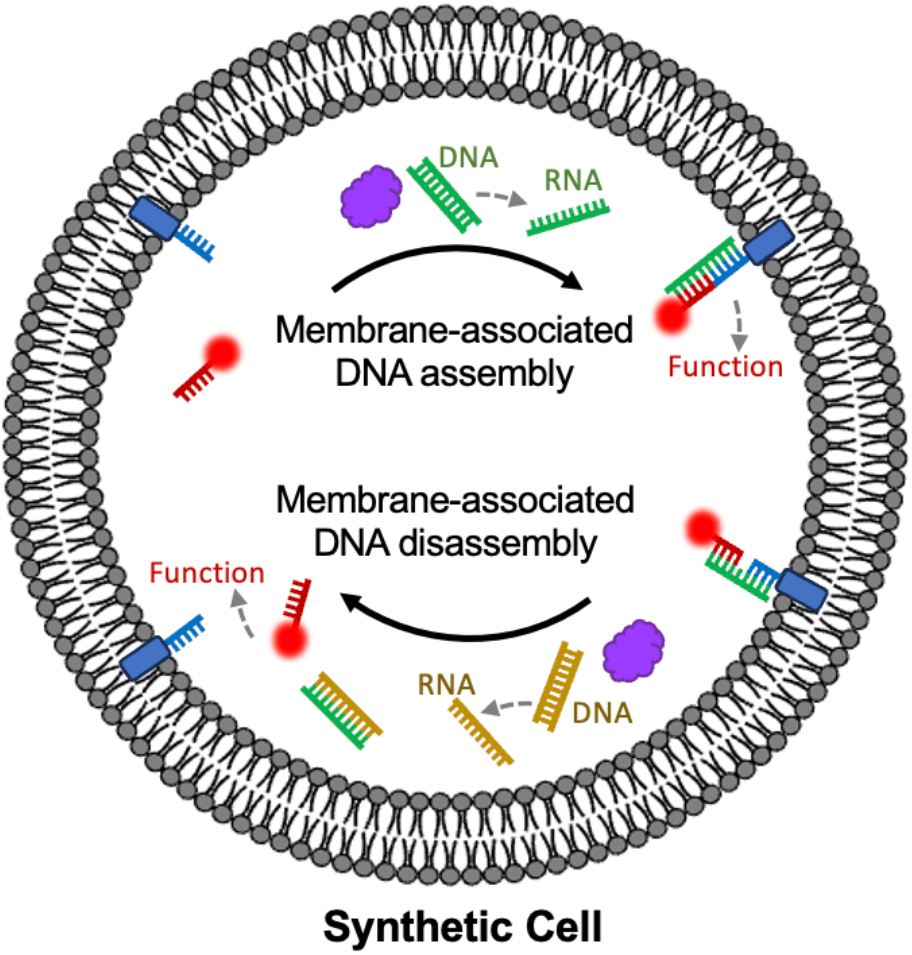

## Introduction

Bottom-up synthetic biology aims to construct minimal cell-like systems from non-living components, providing platforms to study biological processes^1,2^ and engineer programmable biomolecular devices.^3–7^ Giant unilamellar vesicles (GUVs) encapsulating *in vitro* transcription-translation systems have emerged as widely used synthetic cell models, enabling controlled gene expression within cell-like compartmentalised environments.^6–10^ However, although the production of RNA and proteins in synthetic cells is now well established, achieving spatiotemporal control over their organisation remains challenging.^11^

Spatial organisation underpins many biological processes, particularly membrane-associated signalling and molecular clustering (**Figure 1A**).^2^ For example, in living cells, membrane-associated clusters mediate cell-cell adhesion,^12^ membrane fusion and communication,^13^ and cell division.^14^ In contrast, synthetic cell systems are often spatially homogenous, limiting the construction of higher-order behaviours. Existing strategies for membrane localisation typically rely on external inputs or irreversible mechanisms, such as photosensitive anchors or enzymatic cleavage,^15,16^ restricting *in situ* dynamic and programmable regulation of membrane association within synthetic cells.^11^

**Figure 1.**
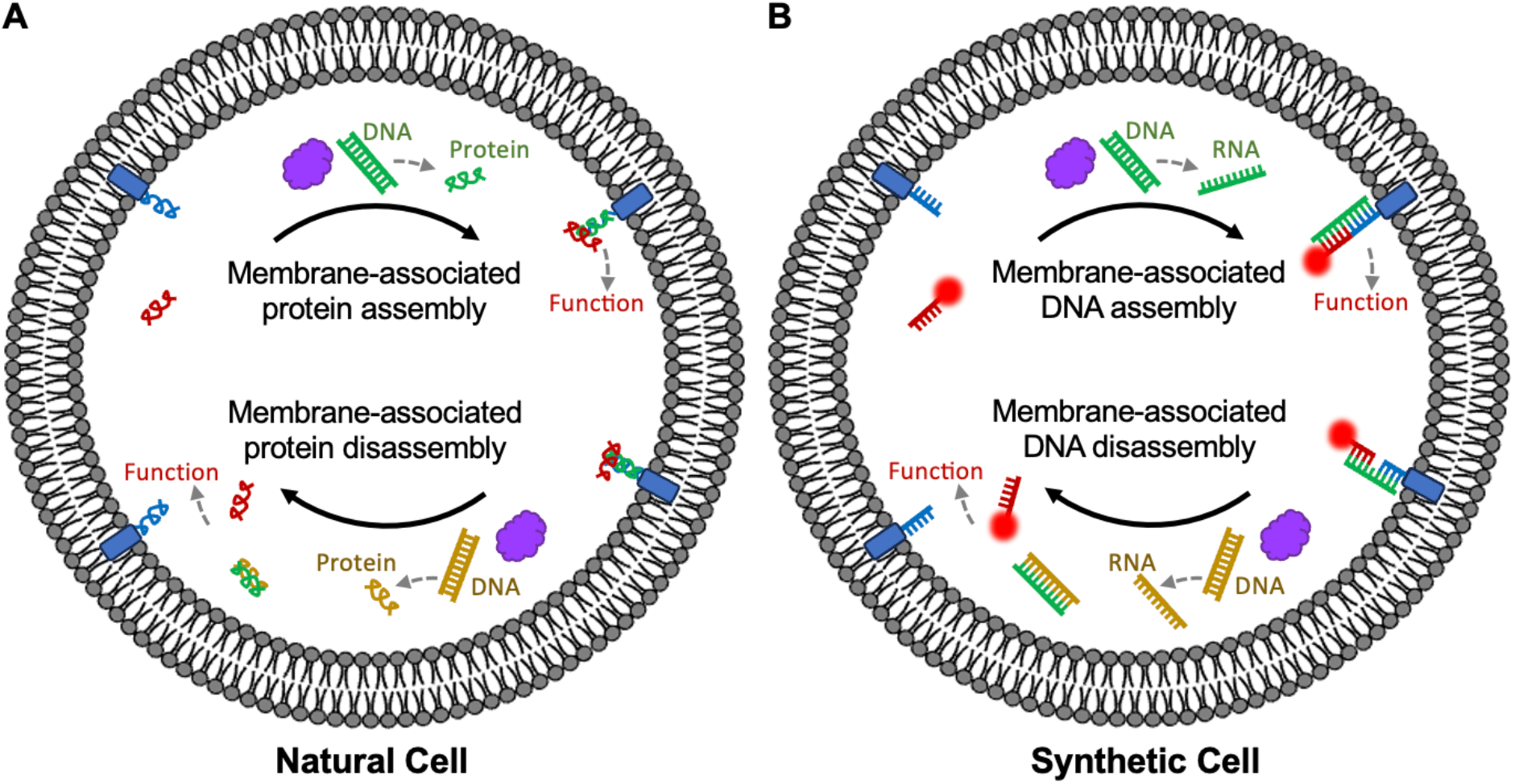
Dynamic membrane organisation in natural and synthetic cells. **A)** In natural cells, gene expression drives the synthesis of proteins that assemble into dynamic membrane-associated complexes to mediate diverse cellular functions. **B)** Analogously, we now show that gene expression can drive programmable and dynamic membrane organisation, in GUV-based synthetic cells, through transcription-controlled DNA assembly. Transcription-generated RNA linkers recruit fluorescent DNA cargo to cholesterol-modified DNA anchors at the membrane. Membrane localisation can be reversed by enzymatic RNA degradation or by transcription-triggered toehold-mediated strand displacement, providing a foundation for engineering genetically-encoded higher-order functions.

DNA nanotechnology provides a powerful framework for programmable molecular organisation through predictable base-pairing interactions and has been widely used to control the assembly and localisation of biomolecules.^17–21^ In parallel, RNA generated by *in vitro* transcription (IVT) offers a dynamic regulatory layer that can couple molecular organisation to gene expression.^9,10^ However, integrating transcriptional processes with DNA nanotechnology inside confined cell-like environments remains non-trivial. Previous studies have shown that enzymatic components, including RNA polymerases, can wreak havoc on nucleic acid systems through non-specific electrostatic interactions.^22–25^ Such interactions can sequester nucleic acid strands, impacting system efficiency, and lead to promiscuous promoter-independent transcriptional activity, limiting transcriptional performance and nanostructure function.

Here, we have managed to combine DNA nanotechnology with IVT, inside GUV-based synthetic cells, to programmably and reversibly control membrane-associated DNA localisation (**Figure 1B**). RNA strands transcribed within the enclosed GUV lumen acted as linkers that recruited fluorescently-labelled DNA cargo to cholesterol-modified DNA anchored at the lipid membrane. The reversal of this membrane localisation could be timed via RNase H-mediated degradation of the RNA linker. Additionally, when using a DNA linker, transcribed RNA could be used to reverse membrane binding via toehold-mediated strand displacement (TMSD). We found that successful integration of transcriptional machinery with nucleic acid nanotechnology requires careful control of reaction conditions and molecular interactions, and that addressing these constraints enables robust operation of the combined system. This work establishes a gene expression-driven platform for programmable and dynamic control of membrane-associated components in synthetic cells, providing a foundation for engineering signalling assemblies and higher-order cellular behaviours.

## Results

### IVT conditions compatible with DNA nanotechnology in GUVs

We first sought to establish IVT conditions compatible with both DNA nanotechnology and stable GUV encapsulation. Commercial IVT systems are typically optimised for bulk reactions to maximise RNA production, but these conditions can disrupt membrane integrity,^26^ cause aggregation,^27–29^ and interfere with nucleic acid assemblies.^23–25,30^ Using a previously reported GUV-optimised *in vitro* transcription-translation buffer,^31^ we developed IVT conditions that supported both tuneable transcriptional activity and robust GUV formation. Components required for translation were omitted and the concentrations of magnesium and ribonucleotide triphosphates (NTPs) were optimised to minimise GUV aggregation. We measured bulk IVT expression of a Broccoli aptamer using this buffer by fluorescent spectroscopy. Surprisingly, the IVT proved highly sensitive to reaction preparation, with RNA yield dependent on the order of component addition during assembly of the transcription mixture (**Supplementary Figure 1**). To our knowledge, this has not been previously reported.

The optimised IVT mix was then encapsulated within GUVs. The GUVs were prepared with egg phosphatidyl choline lipid, using the phase transfer emulsion method.^6,7^ We observed that GUV integrity was adversely affected by use of a commercial T7 RNA polymerase preparation (**Supplementary Figure 2**). Glycerol, which is commonly present in enzyme storage buffers,^32^ is known to perturb lipid membrane stability^26^ and may also influence system function by promoting transbilayer redistribution of cholesterol-modified DNA.^33^ Therefore, we recombinantly expressed and purified T7 RNA polymerase, enabling us to improve the yield of GUVs through lowering the glycerol concentration in the IVT reaction (**Supplementary Figure 2**). IVT functionality in GUVs was confirmed by measuring Broccoli aptamer expression via fluorescence microscopy (**Supplementary Figure 3**). These results demonstrate that careful optimisation of reaction composition and preparation is essential for robust integration of IVT with membrane-based nucleic acid systems.

### IVT-mediated DNA localisation

In order to localise internally encapsulated DNA to the inner GUV membrane, we then designed RNA linkers that could hybridise to two non-complementary DNA constructs (**Figure 2A**). The two non-complementary DNA strands were a target fluorescently-labelled strand used to track localisation (a-DNA), and a cholesterol-modified strand anchored to the inner GUV membrane (b-DNA). These two DNA strands would then be encapsulated in GUVs, along with the IVT system and a DNA template encoding a c-RNA linker. Following *in situ* transcription of the c-RNA linker strand, the a- and b-DNAs become bridged, resulting in membrane localisation of the fluorescent a-DNA. Nucleic acid sequences were designed using NUPACK^34^ to minimise secondary structure formation while incorporating the required 5’-GGG initiation motif for efficient T7 transcription.^35,36^ Sequence specificity was experimentally validated by native polyacrylamide gel electrophoresis (PAGE) (**Supplementary Figure 4A**). The binding behaviour of the cholesterol- and fluorophore-modified constructs was similarly confirmed using native PAGE (**Supplementary Figure 4B**).

**Figure 2.**
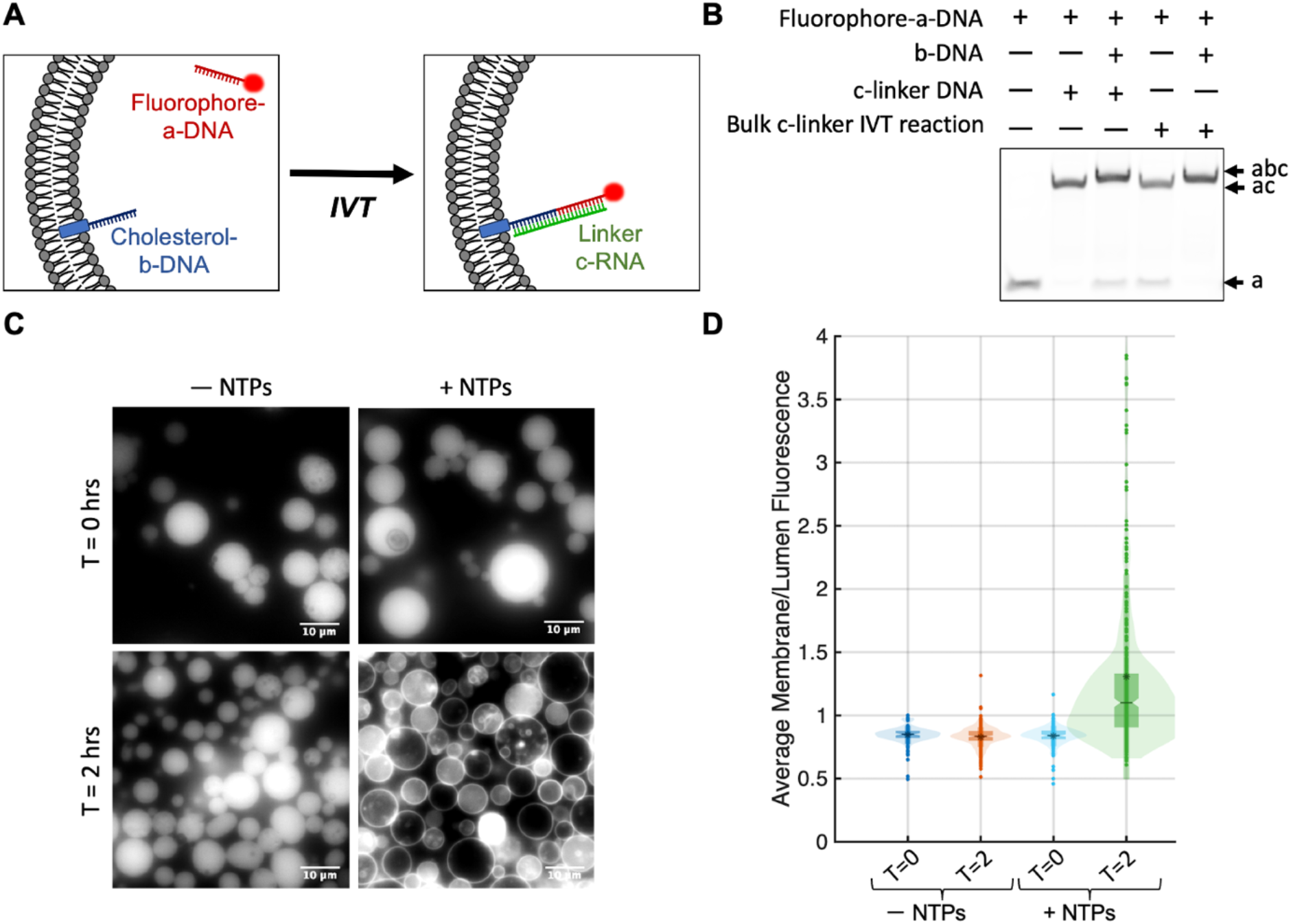
IVT-mediated localisation of DNA to internal GUV membranes. **A)** Schematic of c-RNA-mediated bridging between fluorophore-labelled a-DNA and membrane-anchored cholesterol-b-DNA, inside GUVs. **B)** Native PAGE analysis of fluorophore-a-DNA showing that both c-DNA and c-RNA generated by bulk IVT mediate linking between a-DNA and b-DNA. **C)** Fluorescence microscopy images of GUVs encapsulating fluorophore-a-DNA, cholesterol-b-DNA, and c-RNA IVT reaction components in the presence or absence of NTPs. GUVs were imaged immediately after generation and following incubation at 37 ºC for 2 h. Images are representative of three biological replicates. Scale bar, 10 µm. **D)** Quantification of fluorophore-a-DNA membrane localisation in individual GUVs using a circle-detection-based image analysis workflow. Localisation was quantified as the membrane-to-lumen fluorescence intensity ratio. Mean ratios were 0.85 (− NTPs, T = 0 h), 0.83 (− NTPs, T = 2 h), 0.84 (+ NTPs, T = 0 h), and 1.31 (+ NTPs, T = 2 h). Data from three biological replicates were pooled. Boxplots show the interquartile range (box), median (notch), and mean (asterisk).

We first evaluated whether IVT-generated c-RNA could bridge the a- and b-DNA strands in bulk solution, measured by native PAGE (**Figure 2B**). Fluorescent a-DNA and b-DNA were incubated with IVT reaction components and either a DNA template encoding c-RNA or with a DNA analogue (c-DNA) of the linker sequence for 2 h at 37 ºC. IVT-mediated production of c-RNA generated a distinct higher molecular weight band identical to that observed using c-DNA, indicating efficient bridging between the a- and b-DNA strands. This same system was then carried out on the external surface of GUVs. The external solution contained the same reaction components as above, but using the cholesterol-modified b-DNA. Both the c-DNA and the *in situ* transcribed c-RNA successfully induced membrane localisation of fluorescent a-DNA (**Supplementary Figure 5A**), confirming that the system architecture supported the intended behaviour. However, upon encapsulation within GUVs, membrane localisation was not observed for either IVT-generated c-RNA or the c-DNA analogue (**Supplementary Figure 5B**). These results suggested that transcriptional conditions within the confined synthetic cell environment interfered with DNA nanostructure function.

T7 RNA polymerase is known to interact non-specifically with nucleic acids through electrostatic interactions between positively charged residues and the polyanionic DNA backbone.^37^ Such interactions can reduce strand accessibility and promote promoter-independent transcriptional activity, thereby disrupting DNA-based systems.^22–25^ We identified that pre-incubation of the polymerase with excess c-RNA template DNA on ice for 2 h before reaction assembly enabled functionality within GUVs, consistent with a reduction in non-specific enzyme-DNA interactions (**Figure 2D-E**). Under these conditions, c-RNA generated by *in situ* IVT mediated membrane recruitment of fluorescent a-DNA inside GUVs, increasing the average membrane-to-lumen fluorescence ratio from 0.84 at T = 0 h to 1.31 after 2 h at 37 ºC. In contrast, control reactions lacking NTPs showed no membrane enrichment over the same period (0.85 at T = 0 h and 0.83 at T = 2 h), confirming that localisation depended on active transcription and RNA-mediated bridging. The effectiveness of the pre-incubation strategy depended on both incubation time and the relative concentrations of polymerase and template DNA, supporting a competitive binding mechanism. Because pre-incubation also reduced IVT yield (**Supplementary Figure 6**), transcription conditions were subsequently re-optimised. These findings demonstrate that controlling interactions between transcriptional machinery and nucleic acid components is essential for integrating DNA nanotechnology within confined synthetic cell systems. This requirement may be more pronounced than in bulk reactions because vesicle encapsulation is known to stabilise intermolecular associations and increase nucleic acid binding activity,^38^ while the prolonged incubation required for GUV synthesis provides additional time for these interactions to occur.

### Unbinding of membrane-bound DNA via toehold-mediated strand displacement

In parallel with membrane binding, it is important to develop systems that can unbind DNA from the membrane. To demonstrate IVT-mediated delocalisation of DNA from GUV membranes, we implemented TMSD to dissociate bridged a- and b-DNAs (**Figure 3A**). A modified c-DNA sequence was generated to include a toehold region (ct-DNA), enabling displacement by a fully complementary invader d-RNA strand, generated through IVT. The single-stranded toehold was positioned centrally, rather than terminally, to minimise potential promoter-independent transcription^22,30^ and reduce the likelihood of interference with any future cargo conjugated to the a-DNA terminus. Because efficient TMSD depends on strand accessibility and relative concentrations,^39^ the concentrations of a- and b-DNA were reduced, compared to previous conditions, to ensure that IVT-generated d-RNA was in excess.

**Figure 3.**
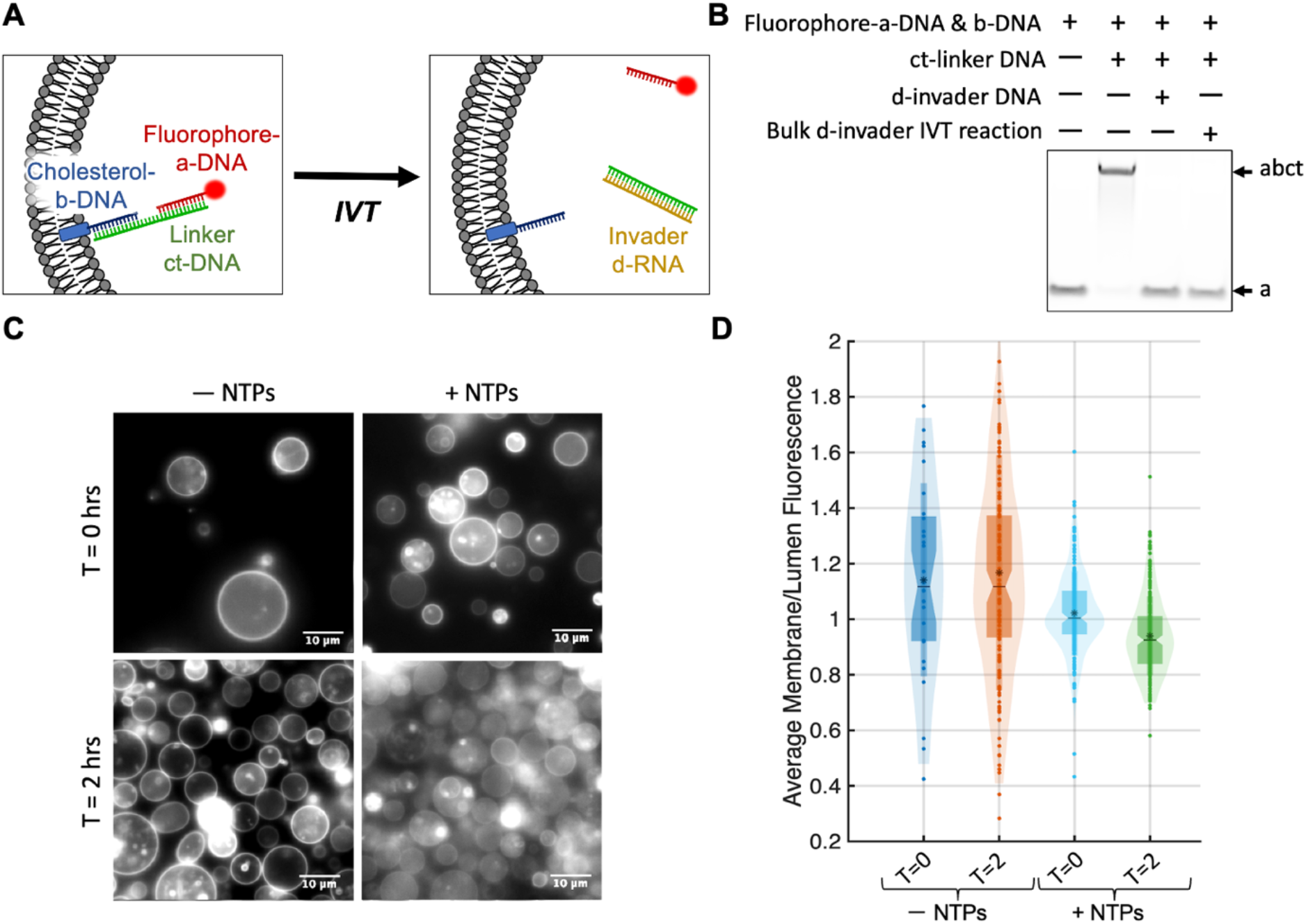
IVT-mediated unbinding of DNA from internal GUV membranes via TMSD. **A)** Schematic of d-RNA-mediated TMSD, resulting in dissociation of fluorophore-labelled a-DNA from membrane-anchored cholesterol-b-DNA. **B)** Native PAGE analysis of fluorophore-a-DNA showing that both d-DNA and d-RNA (generated by bulk IVT) mediate dissociation between the pre-annealed a-/b-/ct-DNA complex. **C)** Fluorescence microscopy images of GUVs encapsulating pre-annealed fluorophore-a-DNA, cholesterol-b-DNA, and linker ct-DNA together with d-RNA IVT reaction components in the presence or absence of NTPs. GUVs were imaged immediately after generation and following incubation at 37 ºC for 2 h. Images are representative of three biological replicates. Scale bar, 10 µm. **D)** Quantification of fluorophore-a-DNA localisation in individual GUVs using a circle-detection-based image analysis workflow. Membrane localisation was quantified as the membrane-to-lumen fluorescence intensity ratio. Mean ratios were 1.14 (− NTPs, T = 0 h), 1.17 (− NTPs, T = 2 h), 1.02 (+ NTPs, T = 0 h), and 0.93 (+ NTPs, T = 2 h). Data from three biological replicates were pooled. Boxplots show the interquartile range (box), median (notch), and mean (asterisk).

We first validated the ability of IVT-generated d-RNA to dissociate the a- and b-DNA strands in bulk using native PAGE (**Figure 3B**). Pre-annealed a-, b-, and ct-DNA complexes were incubated with IVT reaction components and either a DNA template encoding d-RNA or with a DNA analogue invading strand for 2 h at 37 ºC. IVT-mediated production of d-RNA generated a lower molecular weight band corresponding to release of a-DNA from the b-/ct-DNA complex, indicating efficient strand displacement. This system was then transferred into GUVs. Encapsulation of the d-RNA IVT system and cholesterol-b-DNA within GUVs similarly resulted in release of the membrane-localised a-DNA into the vesicle lumen during incubation (**Figure 3C-D**). Generation of the d-RNA reduced the average membrane-to-lumen fluorescence ratio from 1.02 at T = 0 h to 0.93 after 2 h. In contrast, control reactions lacking NTPs retained membrane-enriched fluorescence throughout the same period (1.14 at T = 0 h and 1.17 at T = 2 h), confirming that delocalisation depended on active transcription and the invader d-RNA. Consistent with previous observations, pre-incubation of T7 RNA polymerase with excess template DNA was required for system functionality within GUVs. These results demonstrate that IVT can be coupled to TMSD to programmably reverse membrane localisation of DNA nanostructures in synthetic cells.

### Reversible membrane binding via RNase H-mediated degradation

We next set out to develop a system that could reversibly bind and unbind DNA with temporal control. To achieve this, we co-encapsulated the original c-RNA IVT bridging system with RNase H, inside GUVs (**Figure 4A**). RNase H selectively degrades RNA within RNA/DNA hybrids, providing a mechanism to disrupt c-RNA-mediated bridging between the a- and b-DNAs. Because c-RNA production by IVT initially proceeds rapidly (**Supplementary Figure 6**), tuning the RNase H concentration was expected to enable transient, oscillator-like localisation behaviour.^24,25^ Addition of 2 U RNase H to the encapsulated IVT reaction prevented any membrane enrichment of fluorescent a-DNA (**Figure 4B-C**), consistent with rapid degradation of c-RNA preventing formation of the membrane localised complex. In contrast, addition of 1 U RNase H produced transient localisation. Initially, the average membrane-to-lumen fluorescence ratio increased from 0.82 at T = 0 h to 1.43 after 1 h, demonstrating membrane binding, before decreasing to 0.90 at 2 h, owing to subsequent RNase H-mediated degradation of c-RNA. These results demonstrate that coupling IVT-driven assembly with enzymatic RNA degradation enables temporally-regulated, dynamic membrane localisation within synthetic cells. More broadly, the successful implementation of this behaviour depended on controlled polymerase-DNA interactions, stable IVT activity,^24,25^ and efficient GUV encapsulation; highlighting key requirements for integrating transcriptional networks with DNA nanotechnology in confined cell-like environments.

**Figure 4.**
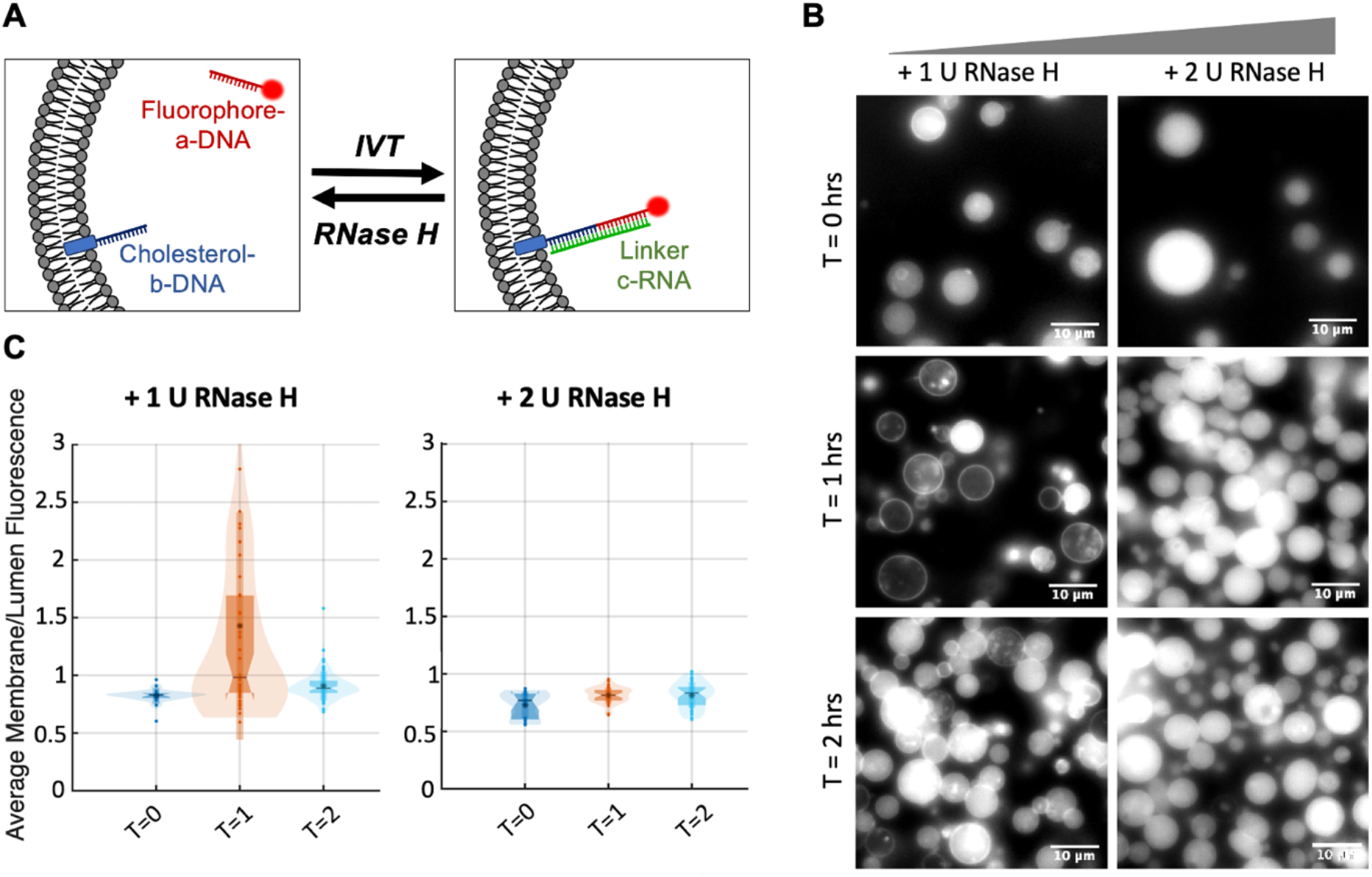
Dynamic localisation of DNA at GUV membranes, mediated by IVT and Rnase H. **A)** Schematic of c-RNA-mediated bridging between fluorophore-labelled a-DNA and membrane-anchored cholesterol-b-DNA, followed by RNase H-mediated degradation of the c-RNA linker. **B)** Fluorescence microscopy images of GUVs encapsulating fluorophore-a-DNA, cholesterol-b-DNA, c-RNA IVT reaction components, and varying concentrations of RNase H. GUVs were imaged immediately after generation and following incubation at 37 ºC for 1 h and 2 h. Images are representative of three biological replicates. Scale bar, 10 µm. **C)** Quantification of fluorophore-a-DNA localisation in individual GUVs using a circle-detection-based image analysis workflow. Membrane localisation was quantified as the membrane-to-lumen fluorescence intensity ratio. Mean ratios for 1 U RNase H were 0.82 (T = 0 h), 1.43 (T = 1 h), and 0.90 (T = 2 h). Mean ratios for 2 U RNase H were 0.73 (T = 0 h), 0.82 (T = 1 h), and 0.82 (T = 2 h). Data from three biological replicates were pooled. Boxplots show the interquartile range (box), median (notch), and mean (asterisk).

## Discussion

We have developed a strategy for gene expression-driven and reversible DNA localisation at synthetic cell membranes by integrating IVT with DNA nanotechnology inside GUVs. RNA strands, transcribed *in situ*, functioned as dynamic molecular regulators that controlled membrane recruitment and release of DNA cargo through hybridisation, strand displacement, and enzymatic degradation. Together, these behaviours establish a framework for coupling gene expression processes to programmable spatial organisation within confined synthetic cell environments.

Although the individual components used here – IVT,^9,10^ DNA nanotechnology,^40–43^ TMSD,^23,30,39^ and RNase H-mediated degradation^24^ – are each well established, their integration inside synthetic cells presented substantial challenges. In particular, transcriptional machinery can interfere with nucleic acid assemblies through non-specific interactions, which sequesters strand accessibility and leads to unintended transcriptional activity.^22–25^ Our results demonstrate that careful control of reaction composition, enzyme preparation, and molecular accessibility is essential for enabling reliable operation of coupled transcriptional and DNA-based systems in confined environments. More broadly, these findings highlight the importance of considering systems-level biochemical compatibility when integrating orthogonal molecular technologies within synthetic cells.^11,44^

The platform presented here provides a route towards increasingly sophisticated spatiotemporal organisation in bottom-up synthetic biology. Because nucleic acid interactions are inherently programmable, the RNA mediator design could readily be adapted to implement more complex regulatory behaviours, including cascaded or orthogonal TMSD circuits,^18^ logic-gated localisation,^30^ or transcription-degradation oscillation.^24^ Similarly, incorporation of light^6^ or magnetically^7^ responsive DNA templates could enable externally triggered and spatially resolved activation of the membrane localisation pathways, providing non-invasive control over synthetic cell behaviour. The ability to control membrane-DNA distribution through phase separation could increase sophistication further,^42,43^ enabling domain-specific localisation.

Beyond fluorescent DNA cargo, the broad chemical modifications possible with nucleic acids offers opportunities to localise diverse biomolecular components to synthetic cell membranes, including proteins,^19,45–47^ polymers,^15,16^ or catalytic motifs.^48–50^ Such capabilities could support controlled molecular clustering, compartmentalised reaction networks, and dynamic signalling assemblies that more closely emulate the spatial organisation found in living systems.^2,12–14^ We anticipate that our integration of gene expression networks with programmable membrane organisation will contribute towards the development of increasingly life-like synthetic cells with emergent and adaptive behaviours.

## Author Contributions

M.J.B. and C.D. conceived the project. C.D. designed, performed and analysed the experiments, with contributions from M.J.B. B.L. and C.N. carried out the T7 RNA polymerase expression and purification. C.D. and M.J.B. wrote the paper.

## Acknowledgements

The authors acknowledge funding from the Biotechnology and Biological Sciences Research Council (BB/W011468/1, BB/T008709/1), Engineering and Physical Sciences Research Council (EP/S022856/1, EP/Y032675/1) and Royal Society (URF\R\231007).

## Data availability

All the data generated in this study are available within the article, the supplementary information, and figures. Source data and code will be made available on Zenodo upon acceptance of the manuscript.

## Supplementary Information

Materials and Methods, DNA sequences, and Supplementary Figs. 1-6.

