## Supplementary Information for "Programmable and Dynamic DNA Localisation at Synthetic Cell Membranes"

### Materials and Methods

#### Materials

All solvents and reagents were purchased from Sigma/Merck unless stated otherwise. T7 RNA polymerase (#M0251), T7 Express *E. coli* (#C2566H), and RNase H (#M0297S) were purchased from New England Biolabs. Standard DNA oligonucleotides were synthesised by Sigma/Merck. Cholesterol- and Alexa647-modified DNA oligonucleotides were synthesised by Integrated DNA Technologies. Egg phosphatidyl choline (Egg PC) was purchased from Avanti Polar Lipids. SecureSeal™ Hybridisation Chambers, Texas Red-dextran (10 kDa), and dimethyl sulfoxide (DMSO) were purchased from Thermo Fisher Scientific. Sucrose and magnesium acetate tetrahydrate were purchased from Fluorochem. Bromophenol blue was purchased from TOCRIS. TEMED (N,N,N',N'-tetramethylethylenediamine) and xylene cyanole were purchased from Bio-Rad. pQE30-His-T7RNAP was a gift from Sebastian Maerkl & Takuya Ueda (Addgene plasmid #124138; http://n2t.net/addgene:124138; RRID:Addgene_124138).[50]

#### Recombinant expression of T7 RNA polymerase

A T7 RNA polymerase construct[50] was transformed into T7 Express *E. coli*, and single colonies were grown in 5 mL Luria-Bertani broth (LB) overnight at 37 °C. Cultures were diluted into 1 L LB and grown at 37 °C to A600 0.6-0.8, then induced with 0.2 mM isopropyl β-D-1-thiogalactopyranoside and expressed overnight at 16 °C. Cells were pelleted (4,000 x g, 20 min), resuspended in lysis buffer (50 mM HEPES pH 7.6, 100 mM NaCl, 5 mM imidazole, 7mM β-mercaptoethanol (BME), 0.1% Triton-X100), and lysed by sonication. Clarified lysate (25,000 x g, 20 min) was applied to a gravity-flow column packed with 2 mL Ni-charged IMAC Sepharose 6 FF (Cytiva), pre-equilibrated in lysis buffer. The column was washed with 15 column volumes of wash buffer (50 mM HEPES pH 7.6, 100 mM NaCl, 30 mM imidazole, 7 mM BME, 0.1% Triton-X100) and eluted with 4 mL elution buffer (50 mM HEPES pH 7.6, 100 mM NaCl, 300 mM imidazole, 7 mM BME, 0.1% Triton-X100). Eluent was concentrated using a 3 kDa MWCO Amicon Ultra-15 filter, exchanged into storage buffer (50 mM Tris-HCl pH 7.9, 100 mM NaCl, 20 mM BME, 50% glycerol, 0.1% Triton-X100), quantified by Nanodrop spectrophotometer, and stored at -80 °C.

#### DNA design and preparation

The a-, b-, c-, ct-, and d-nucleic acid sequences were designed using NUPACK,[33] optimising for specific binding and minimal secondary structure formation. All DNAs were resuspended in water and their concentrations estimated by Nanodrop spectrophotometer. To assess binding specificity, the DNAs were mixed at equimolar concentrations, annealed in 20 mM HEPES (pH 7.6) for 1 h at 37 ºC, then 250 ng of each sample was analysed by native PAGE. Native PAGE gels were made using 16% acrylamide/bisacrylamide (19:1) and 1X TBE buffer, and polymerised using 0.8% ammonium persulfate and 0.05% TEMED. Samples were loaded with a 2X loading dye containing 50% glycerol, 0.01% xylene cyanol, and 0.01% bromophenol blue. Gels were run with 1X TBE buffer at 250 V for 45 minutes, then stained with 3X GelRed® Nucleic Acid Gel Stain and imaged with an Azure 200 Gel Imager (Azure Biosystems).

The a-DNA and b-DNA sequences were purchased with a 5'-terminal Alexa647 fluorescent and 3'-terminal cholesterol modification, respectively. Both modified DNAs were resuspended in water and their concentrations estimated by Nanodrop spectrophotometer and native PAGE (as described above). Double stranded c-RNA- and d-RNA-encoding DNA templates were prepared by mixing equimolar amounts of the leading strand with its reverse complement in 20 mM HEPES (pH 7.6), then annealed by heating to 95 ºC for two minutes before cooling slowly to room temperature.

**Bulk IVT reactions**

To assemble bulk IVT reactions, a 5X master mix solution was prepared consisting of 500 mM HEPES (pH 7.6), 100 mM magnesium acetate, and 7.5 mM spermidine. 10 µL bulk IVT reactions were prepared on ice, using the 5X master mix, with the final reaction containing 100 mM HEPES (pH 7.6), 20 mM magnesium acetate, 1.5 mM spermidine, 280 mM potassium glutamate, and 1.5 mM of each NTP. To this, either 1 µL of commercial T7 RNA polymerase (NEB #M0251) or 0.2 µg recombinantly expressed T7 RNA polymerase was added. For c-RNA and d-RNA expression, where indicated, the polymerase was pre-incubated with 2 µM DNA template on ice for 2 h prior to use and used with double the amount of NTPs. To explore RNA-DNA binding, the relevant DNAs were added to bulk IVT reactions at equimolar concentrations, incubated for 2 h at 37 ºC in a thermocycler, then a 6 µL aliquot analysed by native PAGE. For these gels, rather than staining with 3X GelRed® Nucleic Acid Gel Stain (as described above), Alexa647-a-DNA was imaged using the Cy5 fluorescence setting on an Amersham Typhoon Biomolecular Imager (Cytiva).

For Broccoli aptamer expression, 0.5 µM Broccoli-encoding DNA template and 60 µM DFHBI in DMSO were added to a bulk IVT reaction. These were incubated for 2 h at 37 ºC in a thermocycler. Broccoli-DFHBI fluorescence was measured using a Tecan Spark fluorescence plate reader (Tecan Group) at 455 nm excitation and 506 nm emission wavelengths. For kinetic analysis, samples were incubated in the plate reader at 37 ºC with fluorescence intensity measured every 5 minutes.

**GUV synthesis**

GUVs were synthesised using the inverted emulsion method. For a single GUV condition, 75 µL of Egg PC dissolved in chloroform (25 mg/mL) was transferred to a glass vial. The chloroform was evaporated under gentle $N_2$ flow, whilst the vials were rotated at 45º to distribute lipids evenly up the walls of the vial. Residual chloroform was removed by holding the vial under vacuum for 45 minutes. 0.647 g of 0.22 µM polyethersulfone membrane-filtered mineral oil was added to the Egg PC film to a final concentration of 5 mg/µL Egg PC. The vial was vortexed for 1 minute, tightly sealed using parafilm, then sonicated for 45 minutes at 50 ºC.

GUV outer solution was prepared consisting of 100 mM HEPES (pH 7.6), 20 mM magnesium acetate, 280 mM potassium glutamate, 1.5 mM spermidine, and 200 mM glucose. Bulk IVT reactions were prepared as described above except with the addition of 200 mM sucrose, with the NTPs and relevant DNAs added immediately prior to use as GUV inner solution. 250 µL of 5 mg/mL Egg PC in mineral oil was transferred to 1.5 mL centrifuge tubes, and 100 µL was also layered on top of 200 µL chilled outer solution. 10 µL bulk IVT inner solution was added slowly to the 250 µL lipid-containing oil, ensuring the pipette tip moved constantly to disperse the solution. The tubes were briefly vortexed for 3-5 seconds until cloudy emulsions formed, then added dropwise to the lipid-containing oil previously layered on top of chilled outer solution. These tubes were centrifuged at 16,000 x g for 30 minutes at 4 ºC. The oil phase and outer solution were then discarded, and the remaining pellet resuspended in 10 µL chilled outer solution. The resuspended pellet was transferred to a new tube containing 200 µL chilled outer solution, then centrifuged at 10,000 x g for 10 minutes at 4 ºC. All outer solution was then removed, and the pellet resuspended in 40 µL fresh outer solution.

For c-RNA linking, the IVT inner solution contained 1 µM Alexa647-a-DNA with 2 µM cholesterol-b-DNA, to minimise false negatives when analysed via microscopy, along with 2 µM c-DNA linker control or pre-incubated 2 µM c-RNA-encoding DNA template with 0.2 µg recombinantly expressed T7 RNA polymerase. For external IVT reactions, GUV outer solution was used for the inner solution except for sucrose instead of glucose, and the GUVs were resuspended in a bulk IVT reaction. For d-RNA TMSD, 1 µM pre-annealed Alexa647-a-DNA, cholesterol-b-DNA, and ct-DNA complex and 1 µM d-DNA were used. For RNase H, 0.5 µM Alexa647-a-DNA and 1 µM cholesterol-b-DNA were used with varying volumes of RNase H. The concentrations of a- and b-DNA were reduced for RNase H reactions, compared to initial bridging conditions, to compensate for c-RNA degradation to favour efficient IVT-mediated bridging.

**Imaging & Analysis**

40 µL SecureSeal™ Hybridisation Chambers were attached to microscope coverslips, and 0.1% bovine serum albumin (BSA) in phosphate buffered saline was added, followed by a 2-minute incubation at room temperature. The BSA was removed and chambers were washed twice with GUV outer solution. 40 µL of the sample was pipetted into the chamber and the holes were sealed with double-sided tape. The GUVs were imaged using an IX83 inverted microscope (100 X oil objective lens, GFP and Cy5 filter channels) immediately after generation and following 1 h or 2 h incubation in a 37 ºC oven.

All images per experiment were taken with the same settings, including exposure, brightness, and contrast. Images were auto-captured to minimise photobleaching and processed using ImageJ software (an open-source platform: https://imagej.net/licensing/open-source) with the fluorescence channel brightness normalised across all images within a single experiment. For quantification, all images corresponding to a single sample were saved as individual PNG files. ‘Background’ images were created by manually selecting GUV-free regions of microscopy images (one from each sample) and measuring the mean pixel intensity. PNG files were input into a vesicle analysis script.

**DNA sequences** (lower case text indicates position of the promoter sequence)

| DNA name | 5'-DNA sequence |
|---|---|
| a-DNA | TTCCTGTCCTTCCGTTCCTCCTACTTC |
| Fluorophore-a-DNA | Alexa647-TTCCTGTCCTTCCGTTCCTCCTACTTC |
| b-DNA | CTGATCCCACTGTACCTAACTGTCCTT |
| Cholesterol-b-DNA | CTGATCCCACTGTACCTAACTGTCCTT-TEG-Cholesterol |
| c-DNA | GGGAAGGACAGTTAGGTACAGTGGGATCAGGAAGTAGGAGG AACGGAAGGACAGGAA |
| c-RNA DNA template | TTCCTGTCCTTCCGTTCCTCCTACTTCCTGATCCCACTGTACC TAACTGTCCTTCCCtatagtgagtcgtattaatttc |
| c-RNA DNA template reverse complement | gaaattaatacgactcactataGGGAAGGACAGTTAGGTACAGTGGGA TCAGGAAGTAGGAGGAACGGAAGGACAGGAA |
| ct-DNA | GGGAAGGACAGTTAGGTACAGTGGGATCAGGGAGGAAGCG AGGAAGGAAGTAGGAGGAACGGAAGGACAGGAA |
| d-DNA | GGGCTCTTTGAGCCCTTTTTCCTGTCCTTCCGTTCCTCCTAC TTCCTTCCTCGCTTCCTCCCTGATCCCACTGTACCTAACTGT CCTTCCC |
| d-RNA DNA template | GGGAAGGACAGTTAGGTACAGTGGGATCAGGGAGGAAGCG AGGAAGGAAGTAGGAGGAACGGAAGGACAGGAAAAAGGGC TCAAAGAGCCCtatagtgagtcgtattaatttc |
| d-RNA DNA template reverse complement | gaaattaatacgactcactataGGGCTCTTTGAGCCCTTTTTCCTGTCC TTCCGTTCCTCCTACTTCCTTCCTCGCTTCCTCCCTGATCCC ACTGTACCTAACTGTCCTTCCC |
| Broccoli DNA template | GGGTCTAGGAGCCCACACTCTACTCGACAGATACGAATATCT GGACCCGACCGTCTCCTAGACCCtatagtgagtcgtatta |
| Broccoli DNA template reverse complement | taatacgactcactataGGGTCTAGGAGACGGTCGGGTCCAGATATT CGTATCTGTCGAGTAGAGTGTGGGCTCCTAGACCC |

### Supplementary Figures

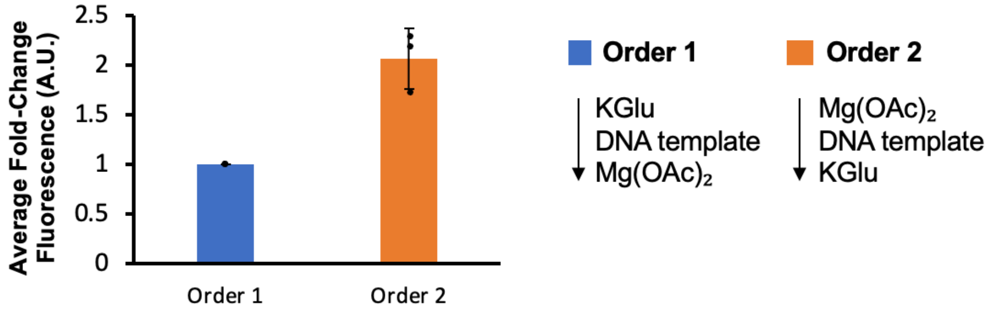

**Supplementary Figure 1. Order of IVT reaction assembly impacts RNA yield.** Bulk IVT reactions containing DFHBI and Broccoli RNA-encoding DNA templates were assembled in varying orders before incubation at 37 °C for 2 h. Fluorescence intensity was used as a proxy for RNA yield. Data are representative of three biological replicates. Values are normalised to 'Order 1' for each biological replicate and are presented as mean ± s.d.

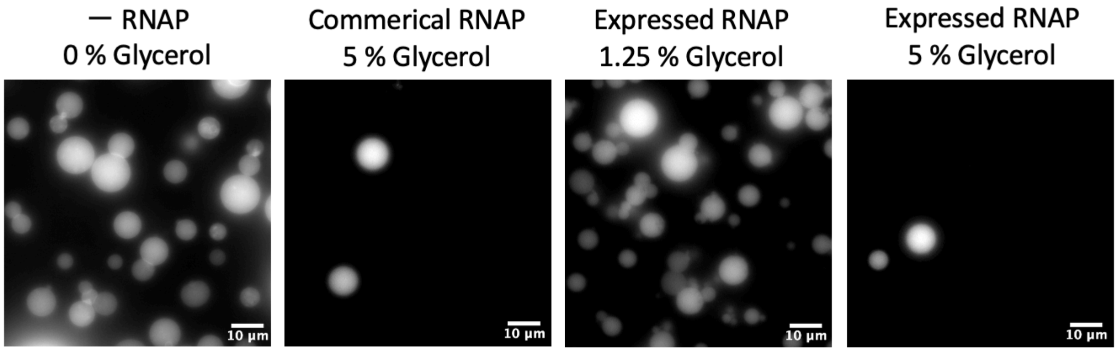

**Supplementary Figure 2. Glycerol reduces GUV yield during encapsulation of T7 RNA polymerase.** Fluorescence microscopy images of IVT GUVs encapsulating Texas Red-dextran together with either no polymerase (RNAP), 1 µL commercially sourced RNAP (NEB #M0251S), or 0.25 µg recombinantly expressed RNAP in the presence or absence of added glycerol. Commercial and recombinant RNAP preparations were suspended in the same storage buffer. Glycerol concentrations refer to the final concentrations within the GUV lumen. GUVs were imaged immediately after generation. Scale bar, 10 µm.

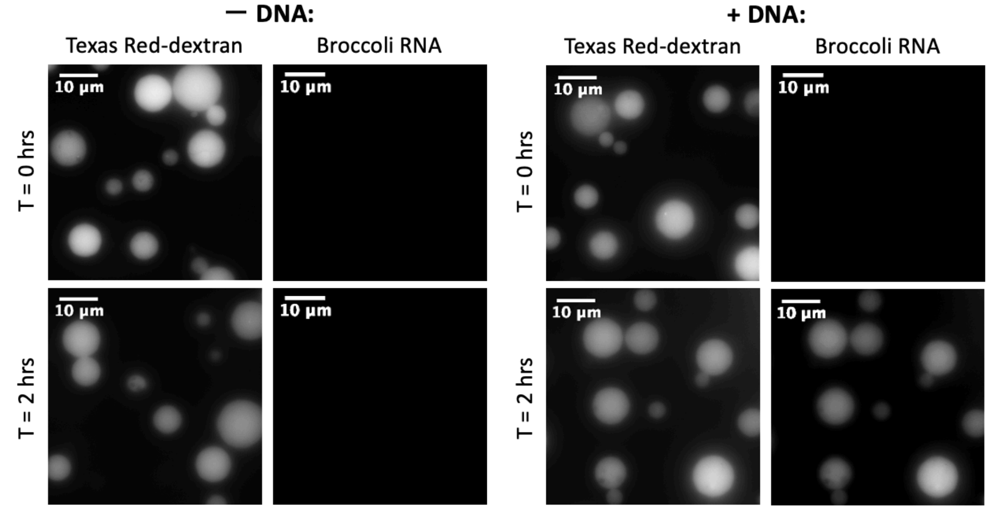

**Supplementary Figure 3. Validation of IVT functionality in GUVs.** Fluorescence microscopy images of IVT GUVs encapsulating Texas Red-dextran dye and DFHBI in the absence (left) or presence (right) of a DNA template encoding for Broccoli RNA. Fluorescence intensity was used as a proxy for RNA yield. GUVs were imaged immediately after generation and following incubation at 37 ºC for 2 h. Scale bar, 10 μm.

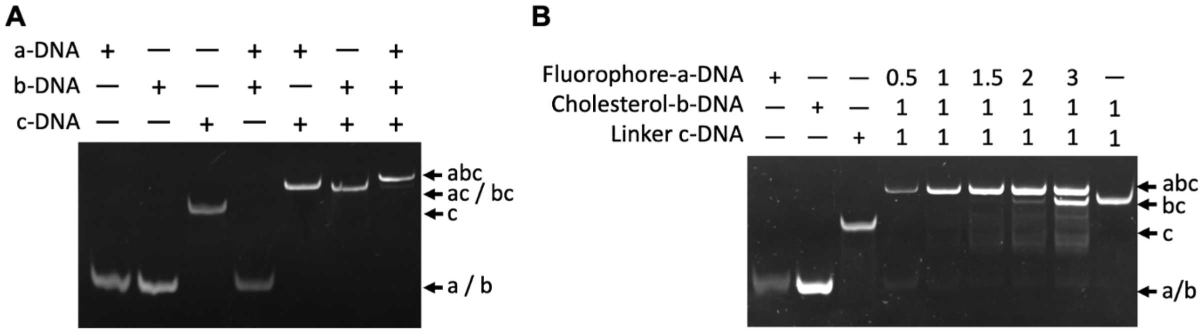

**Supplementary Figure 4. Validation of nucleic acid binding specificity. A)** Native PAGE analysis of a-, b-, and c-DNA strands showing that a-DNA and b-DNA associate only in the presence of linker c-DNA. **B)** Native PAGE analysis of varying molar ratios of fluorophore-labelled a-DNA incubated with cholesterol-b-DNA and c-DNA, demonstrating bridging between the modified DNA constructs.

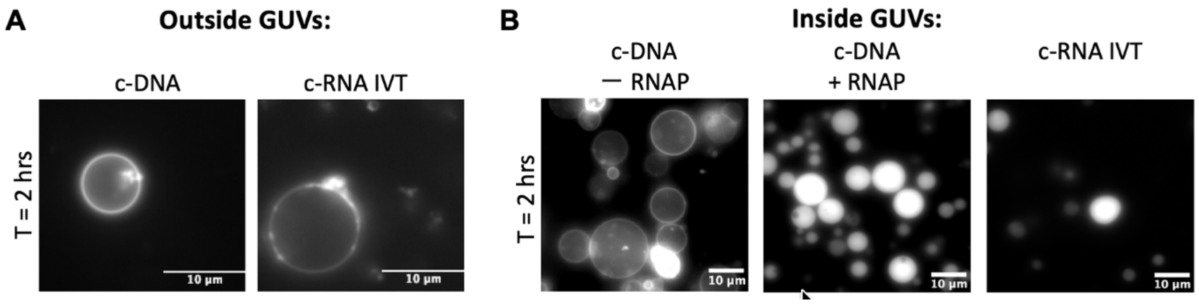

**Supplementary Figure 5. Fluorophore-a-DNA localisation to the GUV membrane was initially only functional when tested externally or in the absence of T7 RNA polymerase. A)** Fluorescence microscopy images of GUVs incubated with external fluorophore-a-DNA, cholesterol-b-DNA, and either c-DNA or c-RNA IVT reaction components for 2 h at 37 ºC. Scale bar, 10 µm. **B)** Fluorescence microscopy images of GUVs encapsulating fluorophore-a-DNA, cholesterol-b-DNA, and either c-DNA in the absence or presence of T7 RNA polymerase and IVT reaction components (RNAP), or c-RNA IVT reaction components. GUVs were imaged following incubation at 37 ºC for 2 h. Scale bar, 10 µm.

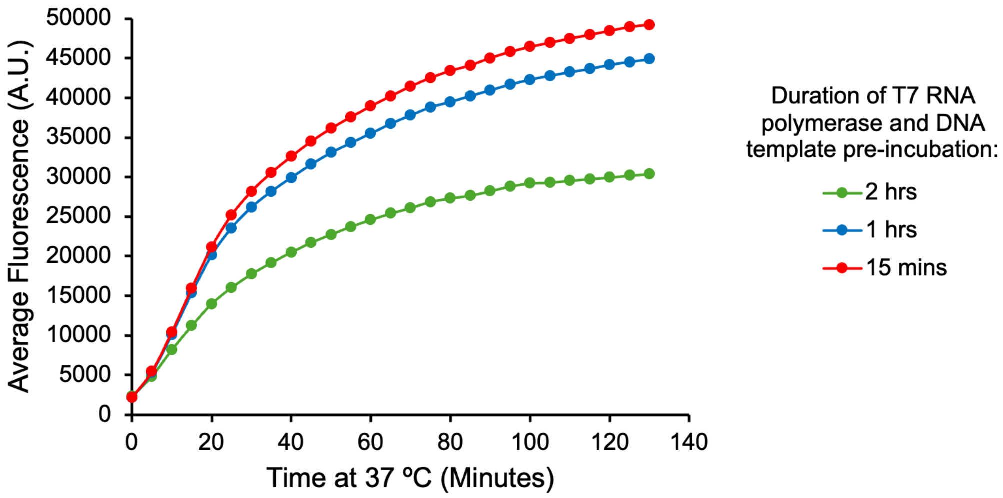

**Supplementary Figure 6. Duration of T7 RNA polymerase and DNA template pre-incubation affects IVT activity.** Recombinantly expressed T7 RNA polymerase (0.2 µg) was pre-incubated with 2 µM Broccoli RNA-encoding DNA template for varying durations before addition to bulk IVT reactions containing DFHBI. Fluorescence intensity was used as a proxy for RNA yield. Data points represent mean values from three biological replicates.
